# Genetic and plastic effects on trait variability in two major tree species: insights from common garden experiments across Europe

**DOI:** 10.1101/2025.07.04.662556

**Authors:** Elisabet Martínez-Sancho, Christian Rellstab, Patrick Fonti, Marta Benito Garzón, Christof Bigler, José Carlos Miranda, Marçal Argelich Ninot, Daniel J. Chmura, Jo Clark, Erik Dahl Kjær, Jon K. Hansen, Manuel Karopka, Mateusz Liziniewicz, Magdalena Nötzli, Aksel Pålsson, Liz Richardson, Evrim A. Şahan, Anne Verstege, Richard Whittet, Yann Vitasse

## Abstract

Phenotypic plasticity and genetic adaptation are key mechanisms that enable species to respond to changing environments. Tree traits do not vary independently but rather in coordination. However, our understanding of whether functional traits are governed by the same mechanism is far from complete. Thus, we aim at assessing the drivers of trait variability of sessile oak and European beech provenances across their distribution ranges.

We estimated growth-related and leaf morphological traits from 9 and 11 provenances of oak and beech, respectively, grown in four different common gardens distributed across their respective distribution areas. Overall, phenotypic plasticity played a dominant role in explaining individual trait variability. For most oak traits, variation among provenances and genetically based plasticity were correlated with the climate of origin, whereas fewer significant associations were found for beech. In oak, climate-transfer distance analyses revealed that traits such as DBH, height, specific leaf area, and long-term growth responses to summer temperature decreased when provenances were moved away from their local precipitation regime. In beech, significant climate-transfer distances were fewer and primarily related to temperature-related parameters. These results suggest that natural selection and local adaptation may play a secondary but notable role. The pattern of multi-trait phenotypes indicates that resource-use strategies among provenances covary with the temperatures of origin in both species.

The limited genetic responses in beech could hinder its survival if it reaches the boundaries of trait plasticity, while oak may better adjust through adaptation. Our study contributes to a better understanding of the interplay between genetic adaptation and phenotypic plasticity in long-lived forest trees.

## INTRODUCTION

Key evolutionary processes, including mutation, genetic drift, gene flow, genetic adaptation and phenotypic plasticity, will determine the ability of tree individuals and provenances to survive under climate change (Burrows *et al*., 2011). Genetic adaptation, driven by natural selection, is the primary evolutionary process that explains how provenances become finely attuned to their local environment (Savolainen *et al*., 2007, 2013). It implies a multigenerational process potentially leading to substantial within-species genetic differentiation over large spatial scales or strong environmental gradients (but see Budde *et al*. 2024). Because trees are sedentary organisms with slow generation turnover (Petit & Hampe, 2006), their adaption pace to rapid climatic changes might be insufficient (Cooper *et al*., 2019; Dauphin *et al*., 2021), potentially enhanced or slowed down by high gene flow that occurs over large distances (Kremer *et al*., 2012). Consequently, tree provenances are often trailing behind their optimal climate (Fréjaville *et al*., 2020), with an increased risk that locally adapted provenances may become maladapted when environmental changes accelerate (Isaac-Renton *et al*., 2018). To persist in a fast-changing environment without compromising their survival, growth, and reproductive capabilities, trees generally exhibit high levels of phenotypic plasticity (Alfaro *et al*., 2014), i.e., the ability of genotypes to express different phenotypes in response to environmental variation (Nicotra *et al*., 2010).

Plant functional traits encompass a set of physiological, morphological and phenological features that determine the individual responses to environmental conditions (Violle *et al*., 2007). The variation in individual traits across species distributions reflects their responses to ecological gradients. The contribution of individual aspects to the observed variation in phenotypic traits can be statistically partitioned into three components: the variation caused by the genotype (G), the variation caused by the environment (E) and the variation caused by the interaction between genotype and environment (G × E). The two latter components contribute to the overall phenotypic plasticity, but only the variation explained by the interaction (genetically based plasticity; e.g., De Jong 1990; Vitasse *et al.,* 2013) quantifies the heritable part of plasticity that differs between genotypes and can thus be subjected to natural selection (Valladares *et al*., 2006; Cooper *et al*., 2019; Fonti *et al*., 2022). The significance of those processes determining intraspecific trait variation differs across species, indicating that species exhibit variation in their evolutionary strategies. Some species show strong evidence of intraspecific genetic variability in trait variation related to climate gradients, pointing to genetic adaptation to the climate of origin, whereas other species more strongly rely on phenotypic plasticity (see Leites & Benito Garzón 2023).

Variation in individual traits cannot be considered in isolation, as it is frequently masked or offset by the influence of other traits, all depending on the overall strategies of the plants (MartínezlVilalta *et al*., 2023). Indeed, trait coordination (Pigliucci 2003; Pigliucci *et al.,* 2006) reflects trade-offs among functions and life-history strategies in response to environmental variation (Benavides *et al*., 2021). Although there is evidence of trait coordination, little is known whether this coordination is maintained across provenances within a species or varies along selective environmental gradients. Traits are theoretically coordinated in spectra aiming to optimize their phenotypes and prioritize some functions following ecological gradients (i.e., leaf economic and wood economics spectra, Chave *et al.,* 2009; Wright *et al.,* 2004; see also the fast-slow plant economics spectrum, Reich 2014). Therefore, trait expression vary along gradients of resource use, dispersal capability, and environmental stress (MartínezlVilalta *et al*., 2023). Since traits covary along the aforementioned axis, plasticity also exists in the manner in which traits covary within each axis of the spectrum (Pigliucci 2003; Rowland *et al.,* 2023). How well these strategies are integrated across a species distribution and their evolutionary drivers are still understudied, yet it would help to better anticipate tree provenance responses to climate change.

Most studies assessing evolutionary processes of trees are based on common garden experiments analysing traits of saplings and early adults (Ramírez-Valiente *et al*., 2021; SolélMedina *et al*., 2022). Although these studies have advanced our understanding in early life-history traits (including survival), they do not reflect the complexity and strategies of older trees, which can vary over time due to ontogenetic or environmental (light/competition) changes or because selection pressure might change. Recent studies exploring dendrochronological time series are emerging and highlight the opportunity to infer a wide range of growth-related traits in adult trees (Housset *et al*., 2018). Tree-ring measurements are a powerful integrative tool to assess tree responses to environmental stresses over time, as any factor limiting or enhancing tree growth is reflected in the size, anatomy, and chemical composition of tree rings (Fritts, 1976). Dendroecological studies have explored climate- growth relations (Babst *et al*., 2019), growth performance (Camarero *et al*., 2015) and/or the effects of extreme climate events (Lloret *et al*., 2011; Anderegg *et al*., 2015). Subsequently, researchers studying provenance genetics and genomics have shown keen interest in these traits and successfully teamed up with dendroecologists in order to, for instance, link growth-derived patterns to natural selection (Heer *et al*., 2018; Avanzi *et al*., 2019) and demographic history (Martínez-Sancho *et al*., 2021).

In this study, we evaluated the role of genetic adaptation and phenotypic plasticity in explaining variation and coordination in phenotypic traits of two major European tree species, sessile oak and European beech. To this end, we collected data from two networks of common garden experiments established in the 1990s and applied dendrochronological techniques to quantify functional and tree-ring-related phenotypes.

Specifically, we addressed the following questions for each study species:

1. What are the contributions of phenotypic plasticity and genetic differences to individual trait variability?
2. Are the differences in trait and multi-trait variation (both genetic and plasticity-based) among provenances associated with their respective climates?
3. Are there signs of local adaptation in individual traits?

## METHODS

### Sampling design

This study investigated two economically and ecologically important European tree species, sessile oak (*Quercus petraea* (Matt.) Liebl) and European beech (*Fagus sylvatica* L.). These two broadleaved tree species from the *Fagaceae* family cover wide distribution areas along large environmental gradients. The study used a network of common gardens established in the 1990s (1990-1993 for oak, see Sáenz-Romero *et al*., 2017; 1998 for beech, see Robson *et al*., 2018). For each species, a subset of four common gardens with contrasting climatic conditions were selected across their area of distribution (Figures 1 and S1, Table S1). For sessile oak, a fully reciprocal subset of nine provenances from the so-called Madsen collection was used (see Lebourgeois *et al.,* 2004). A subset of eleven European beech provenances was used and these were not consistently distributed in all four beech common gardens. Neither common garden network contained an exact local provenance (Table S2).

**Figure 1.**
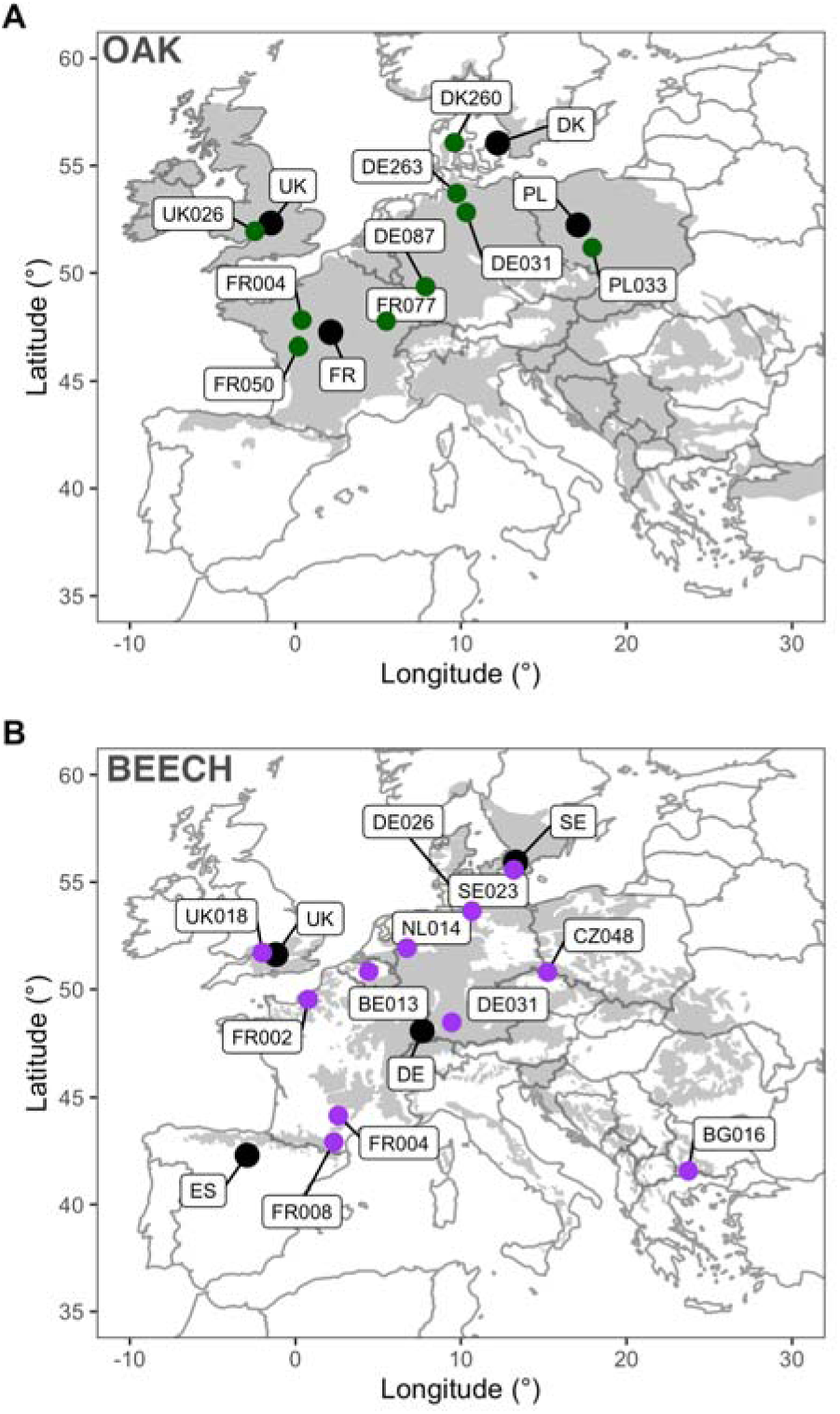
Location of provenances and common gardens of oak (A) and beech (B). Dark grey areas correspond to the species natural distribution ranges obtained from EUFORGEN (www.euforgen.org). Coloured points correspond to the location of the studied provenances (9 for oak and 11 for beech) and the black points refer to the location of the common gardens.

The sampling campaign took place during autumn 2021. For each common garden, sixteen individuals from two blocks (eight trees per block) were sampled at random per provenance. For each individual tree, two increment cores were collected (at 1 m above the ground for maximizing the number of rings) using an increment borer (Haglöf; Långsele, Sweden) attached to a cordless power drill equipped with a torque booster. Tree height and diameter at breast height (DBH at 1.3 m) were recorded using a DBH tape and a Vertex clinometer (Haglöf Vertex 3). The DBH as well as presence/absence of the four contiguous neighbours were also recorded to estimate competition (see below). The Hegyi competition index was calculated based on the DBH of neighbouring trees relative to the DBH of the target tree (Canham *et al*., 2004). Additionally, two leaves were collected from the same individuals. For sessile oak, which commonly hybridizes with other white oak species (e.g., Reutimann *et al.,* 2020), individuals with incorrect species assignment and signs of admixture based on leaf characteristics (Kremer *et al*., 2002; Rellstab *et al*., 2016a) were later discarded.

Details of common garden and provenance locations and abbreviations can be found in Table S1 and Table S2, respectively. Transversal core surfaces were prepared using a sledge microtome (Gärtner & Nievergelt, 2010) to facilitate ring recognition. Images were taken with a digital camera (Canon EOS 5DSR and a 100 mm macro lens, Skippy - https://www.wsl.ch/en/services-produkte/skippy/) at a resolution of 5950 dpi. Ring widths were subsequently measured in CooRecorder v9.6 (Cybis Electronics, Sweden). To estimate the specific leaf area (SLA, mm^2^/mg), three fragments of known area for each leaf were obtained using a punch-borer, dried for 48 hours at 60 °C and their dry mass determined.

### Climatic data

Time series of monthly maximum and minimum temperature as well as total precipitation from the eight common gardens were extracted at 30 arc sec resolution from CHELSA v2.1 (Karger *et al*., 2017) for the period 1987-2021. Similarly, we also extracted time series for all provenances (climate of provenance) from the CHELSAcruts for the climate normal 1961– 1990, which represents the baseline climate prior to seed collection. A total of 19 bioclimatic variables for provenances and common gardens were then calculated for each provenance using the R-package *dismo* (Hijmans *et al*., 2011) including annual mean temperature, mean diurnal range, isothermality, temperature seasonality, maximum temperature of warmest month, minimum temperature of coldest month, temperature annual range, mean temperature of wettest quarter, mean temperature of driest quarter, mean temperature of warmest quarter, mean temperature of coldest quarter, annual precipitation, precipitation of wettest month, precipitation of driest month, precipitation seasonality (coefficient of variation), precipitation of wettest quarter, precipitation of driest quarter, precipitation of warmest quarter and precipitation of coldest quarter.

### Phenotypic traits

Eleven phenotypic traits were investigated, which are divided into five categories (see Table 1 for definitions and abbreviations): i) *tree size* including diameter at breast height (DBH) and height; ii) *leaf morphology* represented by the specific leaf area (SLA); iii) *tree growth increment* including basal area increments (BAI); iii) *long-term growth responses to climate* including the correlation between tree growth and spring and summer mean temperature and precipitation; and iv) *short-term growth responses to extreme drought* including three resilience components.

**Table 1.**
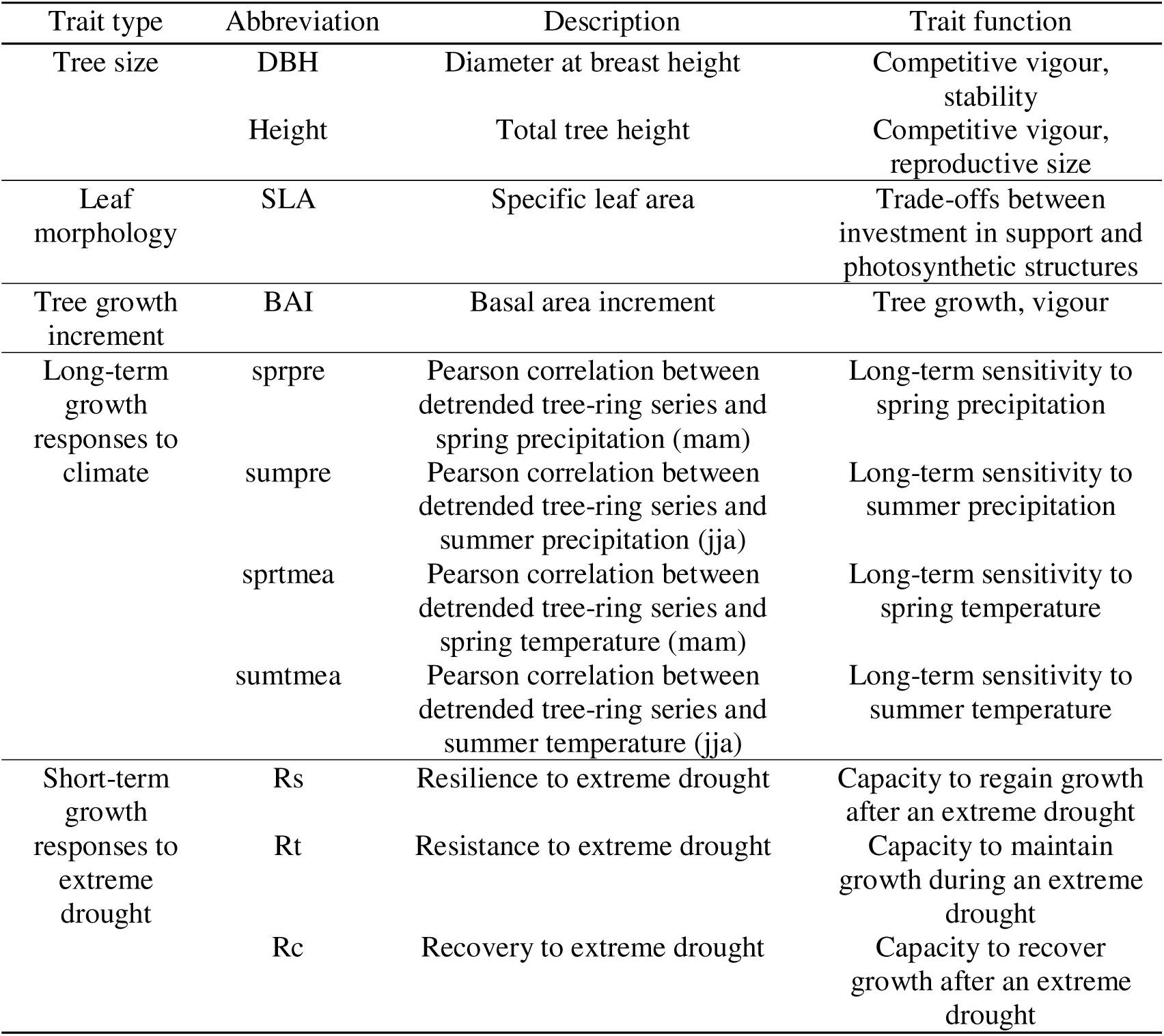
List of phenotypic traits examined and their function.

For tree growth increment, tree-ring series were averaged per individual and then converted into BAI (Biondi & Quedan, 2008). Only BAI from 2006 onwards were considered for the analyses to avoid the juvenile growth of both species. To assess long-term growth responses to climate, first individual tree-ring width series were detrended by fitting a cubic smoothing spline with a 50% frequency cut-off at 20 years using the R-package *dplR* (Bunn, 2010). The obtained detrended series were then correlated with spring (March-April-May) and summer (June-July-August) mean temperature and precipitation for the period 2001-2021 using bootstrapped Pearson correlations from the R-package *treeclim* (Zang, 2016). Short-term growth responses to extreme drought were assessed using the resilience components defined in Lloret *et al*. (2011): resistance (Rt) is defined as the ratio between growth during the drought period and the preceding period, recovery (Rc) as the ratio between growth during the post-drought period and the drought period, and resilience (Rs) as the ratio between growth during the post-drought and pre-drought period. We delimited pre- and post-drought periods to two years to reduce the likelihood that the lag effect of a given extreme climatic event does not overlap with a different extreme event (Anderegg *et al*., 2015). These indices were calculated based on the BAI series. The most extreme drought event was defined based on climatic water balance. The water balance was calculated as the difference between the sum of precipitation and the annual sum of potential evapotranspiration for different time windows (see Figure S2). Potential evapotranspiration was derived from mean temperature using the formula of Thornthwaite from the R-package *spei* (Vicente-Serrano *et al*., 2010). The resulting selected years were 2017 for the beech trial “Pazuengos” (Spain) and the oak trial “Vierzon” (France) and 2018 for the other common gardens (Figure S2). These years correspond to the lowest summer water balance and coincide with the narrowest ring of each site chronology.

### Statistical analyses

All analyses were performed for each species separately. Linear mixed-effects models (Zuur *et al*., 2009) were used to assess the proportion of the variance of the studied traits explained by genetic origin (G), the environment (plasticity, E), and interaction between genetic origin and environment (genetically based plasticity, G × E) using the R-package *lme4* (Bates *et al*., 2015). Individual traits were modelled as a response variable and E (common garden), G (provenance) and G × E (provenance-by-common garden interaction) were treated as fixed effects. Competition was also included as an additive fixed effect whereas block within common garden was considered as a grouping variable for the random intercept. Diagnostic plots were checked for heteroscedasticity and non-normal distribution of the residuals, and log-transformations of the response variable were applied whenever needed. Least square means of provenances, sites and interaction terms were extracted from the respective models using the R-package *lsmeans* (Lenth, 2016). We tested for significance using type III ANOVA for fixed-effects factors, and likelihood ratio tests for random effects. The variance explained by each model was assessed with the R-package *MuMIn* that estimates the pseudo- R^2^, which splits the variance into marginal (R^2^marg, solely explained by the fixed-effects) and conditional (R^2^cond, explained by fixed and random effects) (Nakagawa & Schielzeth, 2013). To understand the importance of G, E and G × E for each of the traits, the marginal variance was then further explored and split into the variance explained by each of the studied fixed factors using semi-partial R^2^ and inclusive R^2^ from the R-package *partR2* (Stoffel *et al*., 2021).

Using these specific mixed-effects models, among-provenance genetic trait variation (genetic effects) was obtained from the least square means of provenances. Among-common garden provenance trait variation (genetically based plasticity) was determined as the maximum range of the genetic effects across common gardens. To explore the potential influence of climate conditions at provenance, both genetic effects and genetically based plasticity of each individual trait were correlated with the 19 bioclimatic variables describing the provenance (1961–1990) using Pearson correlations and controlling for false discovery rate with the Holm method (Holm, 1979).

For each species, climate transfer distances were calculated for each trait as the difference between individual bioclimatic values at the provenance origin (for the period 1961-1990) and the bioclimatic values of the four corresponding common gardens (for the period 1987- 2021). Local climate adaptation was tested for each bioclimatic variable by examining the association between climate transfer distance and mean trait values per provenance and common garden derived from the linear mixed-effect models (effects of G × E from the models) using quadratic functions. We selected potential adaptation signals based on the following criteria: i) significant fitting of the quadratic function (*p* < 0.05), ii) significantly better fit than a linear model (ANOVA, *p* < 0.05), and iii) greater parsimony compared to a linear model (ΔBIC > 2). If more than one bioclimatic parameter met these criteria for a specific trait, the model explaining the higher variance (*R²*) was selected as the best predictor, although both were reported.

The potential influence of climate conditions at provenance on the multi-trait provenance- phenotype was also assessed. To combine multiple traits into a multi-trait phenotype, principal component analysis (PCA) was performed on the least square means of provenances (only G effects obtained from the linear mixed-effects models) of the studied traits. Provenance trait means were scaled and centred before performing the PCA. Linear regressions between the provenance scores along the first two axes of the PCA and the 19 bioclimatic variables of the climate of origin of the provenances were calculated to identify the significant climatic drivers.

The plasticity of the multi-trait phenotypes was quantified in a multivariate framework following Collyer & Adams (2007; SolélMedina *et al*. (2022) and Rowland *et al*. (2023). For each species, a PCA containing the provenance trait means for each provenance at each common garden (G × E effects obtained from the linear mixed-effects models) was calculated. The pairwise trajectories of each multi-trait provenance-phenotype among sites were then determined and the magnitude and direction of change were calculated on the first two PCs. The trajectory magnitude (distance) describes the quantitative difference in multi-trait mean values among sites for each provenance whereas the vector direction (calculated as its angle) represents changes in trait covariation between sites for each provenance (Collyer & Adams, 2007). The maximum magnitude and direction were estimated per provenance and were also correlated with the 19 bioclimatic variables of the climate of origin of the provenances.

All statistical analyses were performed in the R environment (version 4.1.2, R Development Core Team, 2021) and the figures were created using the R-package *ggplot2* (Wickham, 2009).

## RESULTS

### Trait variation among provenances and common gardens

The variation in response to common garden conditions (E) played a variable but mostly dominant role in explaining the non-residual variance of the oak traits, ranging from 5-58% (Figure 2). Specifically, common garden accounted for the largest proportion of variance in specific leaf area (SLA), spring precipitation-growth correlations (sprpre), summer temperature-growth correlations (sumtmea), resilience (Rs), resistance (Rt), and recovery (Rc) to extreme drought. Competition had a significant effect on most oak traits explaining a large part of DBH, basal area increments (BAI) and height. In oak, the interaction between provenance and common garden (G × E, 0-16%) explained slightly more variance in traits than provenance alone (G, 0-7%) and had a significant effect on all traits, with height, DBH, resilience to extreme drought (Rs) and Rc showing the highest variance explained by it. In contrast, provenance (G) significantly influenced height, DBH, BAI, spring and summer temperature-growth correlations (sprtmea and sumtmea), with height exhibiting the highest proportion of variance explained by provenance (7%).

**Figure 2.**
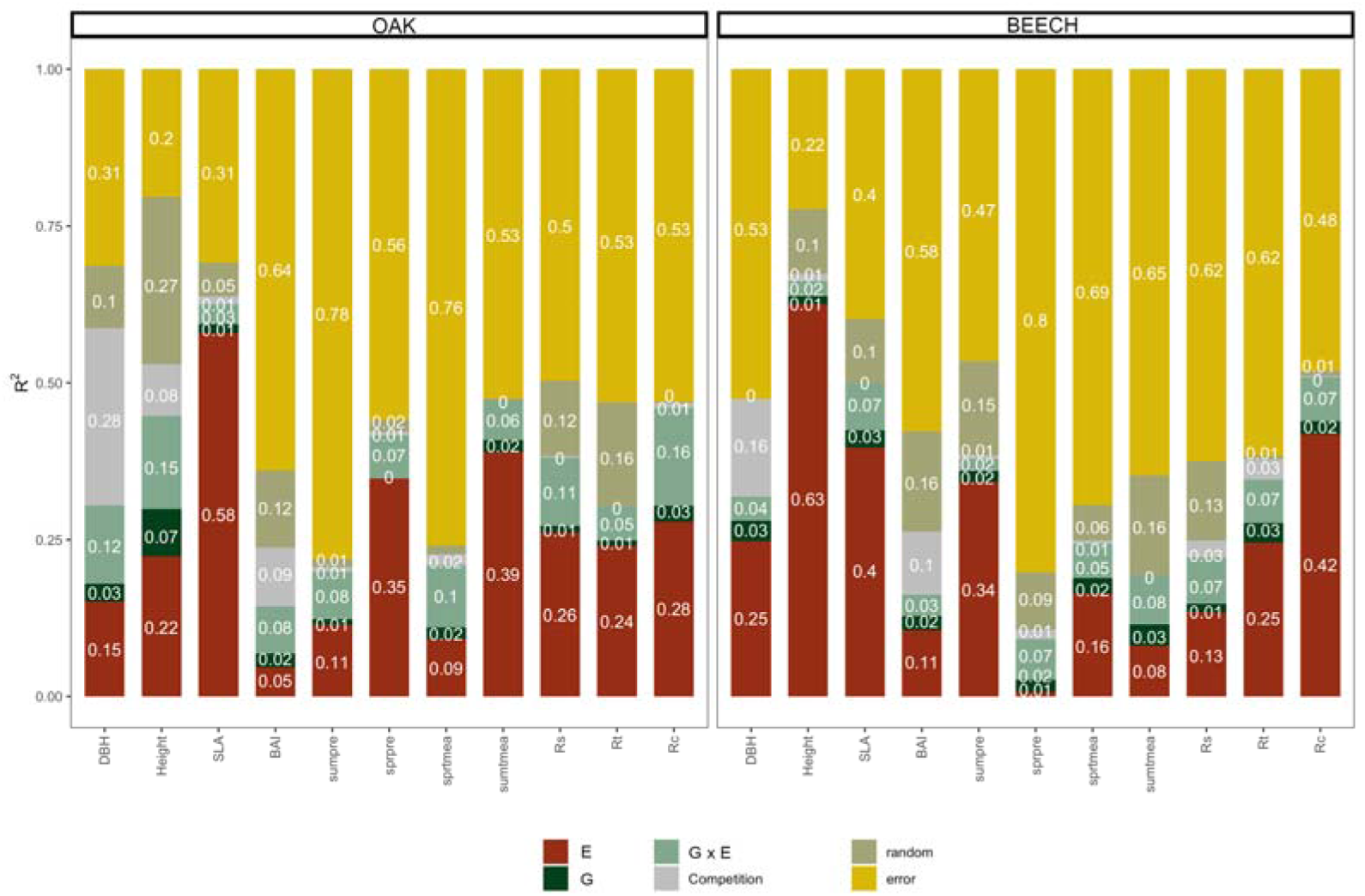
Variance explained (R^2^) by genetic and environmental components of all studied traits. E, proportion of the variance due to common garden; G, proportion of the variance due to provenance; G × E, proportion of the variance due to the interaction between provenance and common garden. The proportions were extracted from the linear mixed-effect models reported in Table S3. Abbreviations are listed in Table 1.

For beech, height, SLA, sumpre, and Rc had more than 30% of their variance explained by common garden (E), while all traits exhibited a significant effect (Table S3). The provenance-by-common garden interaction (G × E) explained less than 10% of variance in all the traits (Figure 2). As in the case of oak, competition had a significant role in most of the trait variability except for sumtmea and Rc. In this species, provenance (G) did not significantly affect sprpre and sumpre, with DBH, SLA, sumtmea and Rt being the traits with the highest proportion of variance explained by provenance.

### Correlation between trait variation and climate of origin and common garden

The least square means of traits from oak provenances (G effects) showed significant correlations mostly with temperature-related variables of the provenances (Figure 3A). Overall, provenances from warmer sites (higher mean annual temperatures, mean temperature during the driest quarter, and/or minimum temperatures during the coldest month or quarter) exhibited smaller heights, weaker sprpre, but stronger sprtmea and higher Rs. Isothermality also correlated significantly with sprtmea, Rs and Rt. Provenances from areas with warm temperatures during the wettest quarter showed larger DBH, BAI and height. Also, oaks from provenances with higher precipitation during the warmest quarter exhibited larger BAI and height, and stronger correlations with sprpre. Provenances with high precipitation level during driest and wettest quarter displayed high sumpre and sumtmea, respectively. Precipitation seasonality at the seed provenance also influenced sumpre and sumtmea.

**Figure 3.**
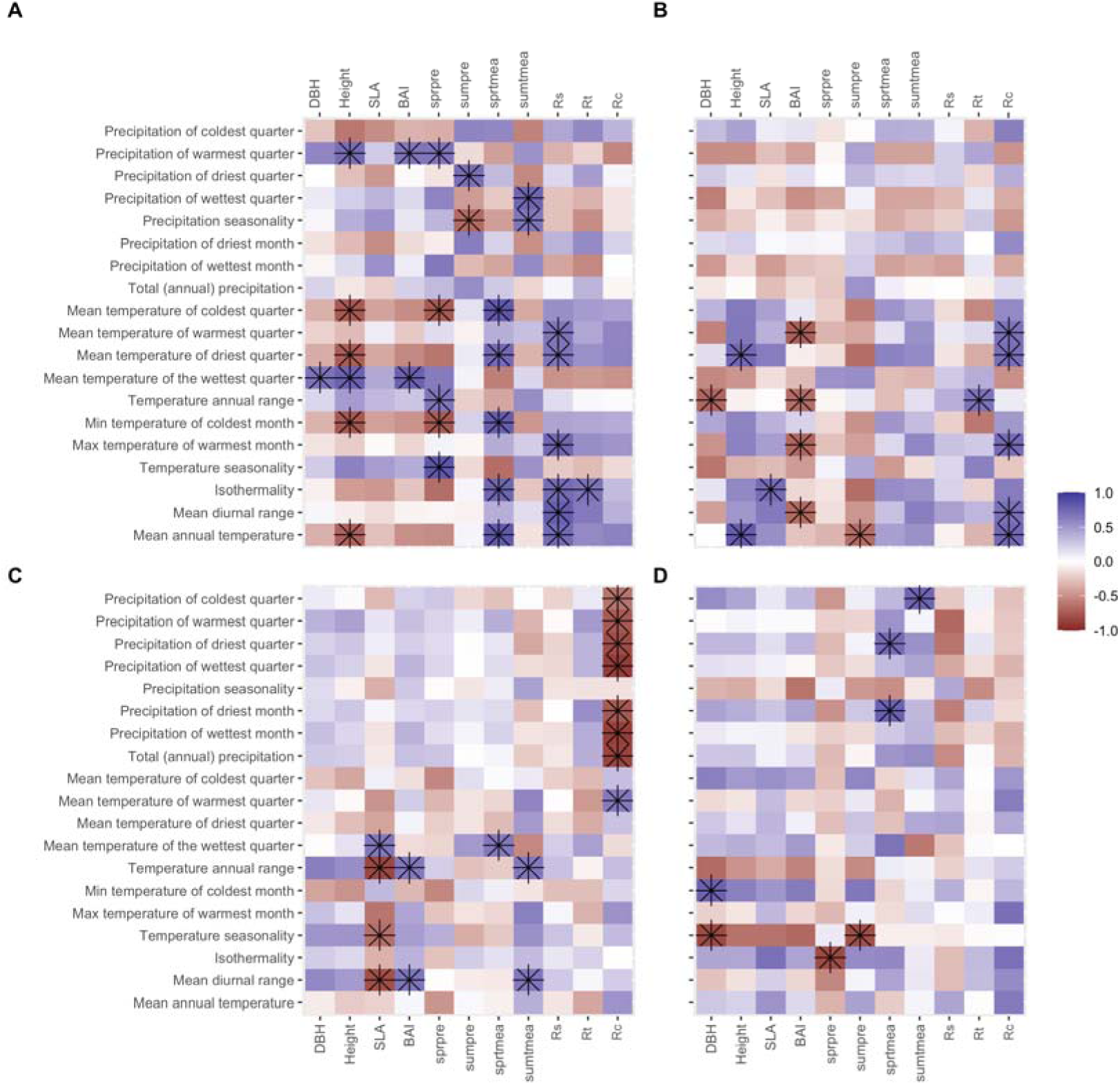
Correlations between traits and climate of origin (1961–1990). Pearson correlations between climate of seed provenance with the mean provenance per trait (A, C), and their range across common gardens (B, D). The upper panels (A, B) refer to oak and the lower ones to beech (C, D). Abbreviations are listed in Table 1. Asterisks indicate significant correlations at *p*< 0.05 level.

The variables related to temperature of the provenance were the primary variables driving the interaction of provenance and common garden (G × E) in oak traits (Figure 3B). Oaks originating from warmer locations exhibited a greater capacity for change in height and Rc but showed a lower change in BAI and sumpre across common gardens. Temperature ranges (either diurnal or annual) and isothermality were also significantly correlated with changes in SLA, BAI, Rc, and Rt.

Beech exhibited fewer significant correlations between least square means of traits (G) and the climate of their provenance (Figure 3C). Beech trees originating from regions with high diurnal or annual temperature range exhibited lower SLA but larger BAI and stronger sumtmea. SLA and sprtmea were also higher for provenances from sites with higher temperature during the wettest quarter. Rc showed significant negative correlations with precipitation-related variables from the provenance. Few correlations were observed between climate of origin and the interaction of provenance and common garden of beech traits.

Temperature seasonality and isothermality were negatively associated with the capacity for change in DBH and with sumpre and sprpre, respectively. Provenances from areas with high precipitation during the driest month/quarter and the coldest month exhibited a higher capacity for change in sprtmea and sumtmea, respectively (Figure 3D).

We also explored the effect of climatic conditions at the common gardens (E) on traits. Oak traits, namely height, SLA, sprtmea and sumtmea, sprpre, and Rt correlated significantly with temperature conditions of the common gardens whereas Rc was significantly associated with precipitation-related variables (Figure S3A). Beech traits, namely SLA, sumpre and Rs, were correlated with precipitation of the common gardens (Figure S3B) whereas DBH, sprpre, sprtmea, Rt and Rc were associated with temperature-related variables.

### Climate-transfer functions

Climate-transfer functions revealed differences between species in terms of adaptation patterns (Figure 4). For oak, height and sumtmea decreased when provenances were transferred away from their local precipitation regime i.e., to more moist sites (bio12) (negative climatic transfer values on x-axis of Fig. 4) or to drier sites (positive values in Fig 4). Similarly, we found a strong relationship between annual precipitation (bio12) and SLA, DBH, and BAI. However, SLA showed a more pronounced decrease when provenances were moved to wetter conditions, while DBH and BAI exhibited stronger declines under drier conditions. Local climatic adaptation patterns in sumpre were also observed when provenances were moved away to a different diurnal temperature range (bio2) and in Rc when moved away to areas with different precipitation seasonality (bio15). DBH, BAI and sumtmea were also significantly related to isothermality but to a lesser degree (bio3) (Table S4). The maximum aligns relatively well with the 0 value in most of the traits, except for SLA and Rc where a shift towards positive values was observed.

**Figure 4.**
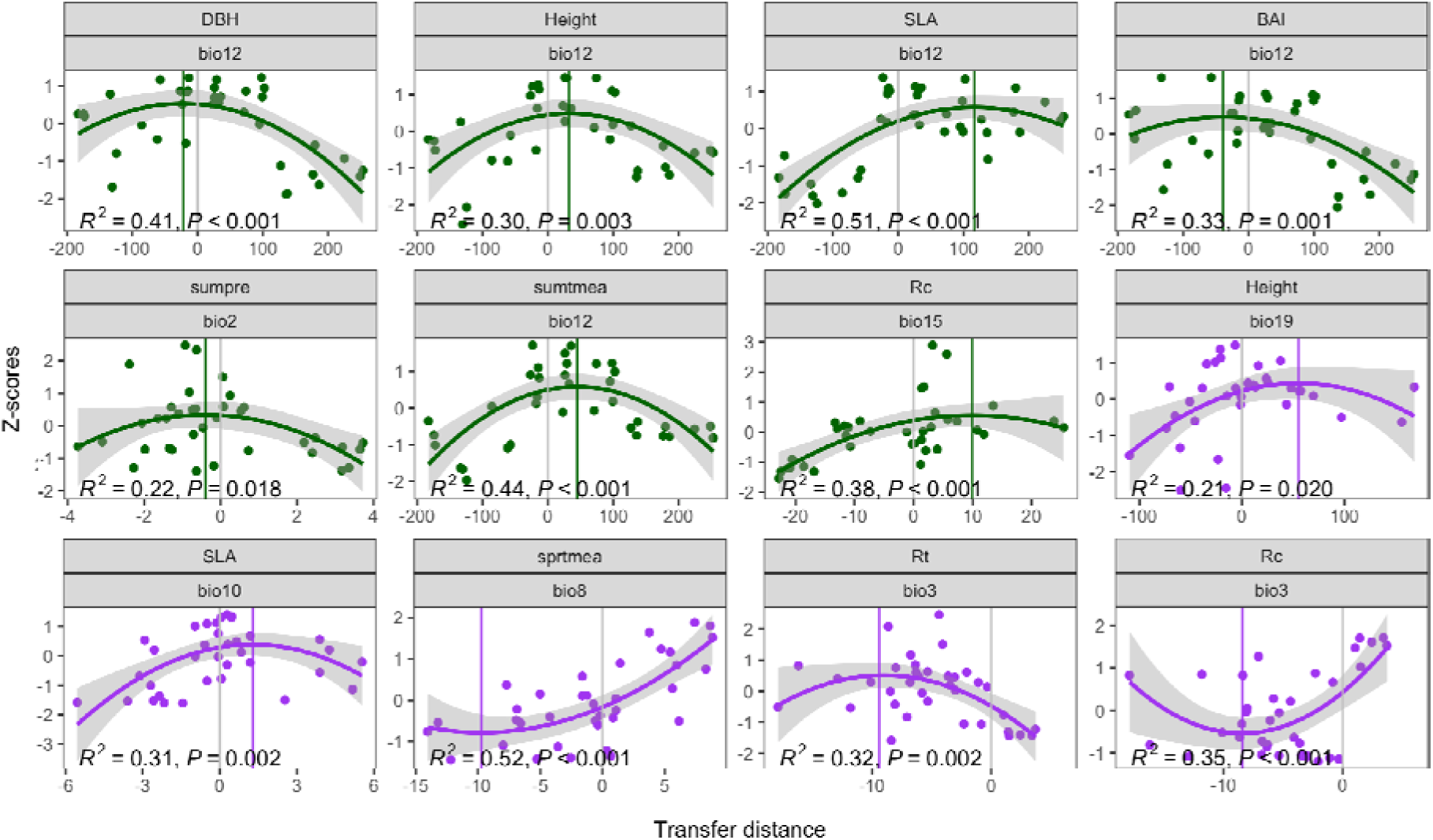
Climate-transfer distance analyses. Fitted significant quadratic functions between the climate transfer distance for each bioclimatic parameters and mean traits by provenance at each common garden for each trait (effects of G × E of the linear models). The climate transfer distance is calculated as the difference between individual bioclimatic values at the provenance origin (for the period 1961-1990) and the bioclimatic values of the four corresponding common gardens (for the period 1987-2021). The green dots and lines are related to oak, while the purple ones correspond to beech. Green and purple vertical lines correspond to the optimum value predicted by the quadratic function for the specific trait. Trait data were normalized (z-scored) for comparison. Bio2, mean diurnal range; bio3, isothermality; bio8, mean temperature of the wettest quarter; bio10, mean temperature of warmest quarter; bio12, total annual precipitation; bio15, precipitation seasonality; bio19, precipitation of coldest quarter.

In contrast to oak, we observed a more varied set of drivers for local climatic adaptation patterns in beech (Figure 4). Height significantly decreased when provenances were moved away from their local precipitation range, i.e., to areas where precipitation during the coldest quarter (bio19) was either reduced or increased. For SLA, this pattern was strongly associated with the temperature of the warmest quarter and month (bio10 and bio5). Sprtmea increased when provenances were moved away from the local climate to areas with a colder wettest quarter (bio8), but the same effect was not observed when provenances were moved to warmer areas. Rt and Rc exhibited opposing patterns with respect to isothermality (bio3); Rt decreased while Rc increased when provenances were moved away from the vertex of the quadratic function. Notably, we observed that in most traits (to a lesser degree in SLA), these maximum (and minimum for Rc) values were shifted from 0.

### Multi-traits and correlations with climate of origin

The PCAs conducted using the least square means of traits (G) revealed a different pattern of trait variation for the two species (Figure 5). For oak, PC1 (38.6% of variance explained) was negatively correlated with height, SLA, BAI, DBH, sprpre and sumtmea, while showing a positive correlation with Rc, Rs, and Rt, and sprtmea. PC2 (19.6%) was positively correlated with DBH, BAI, sumpre and sprtmea, Rt and Rs, and negatively with sumtmea and sprpre. Linear regressions showed a significant relationship between PC1 scores of the provenances and their mean temperature of the driest quarters at the origin (Figure 5A, inset plot) and also with their mean annual temperature (R^2^= 0.66, *p*-value<0.01). No significant pattern was observed for PC2.

**Figure 5.**
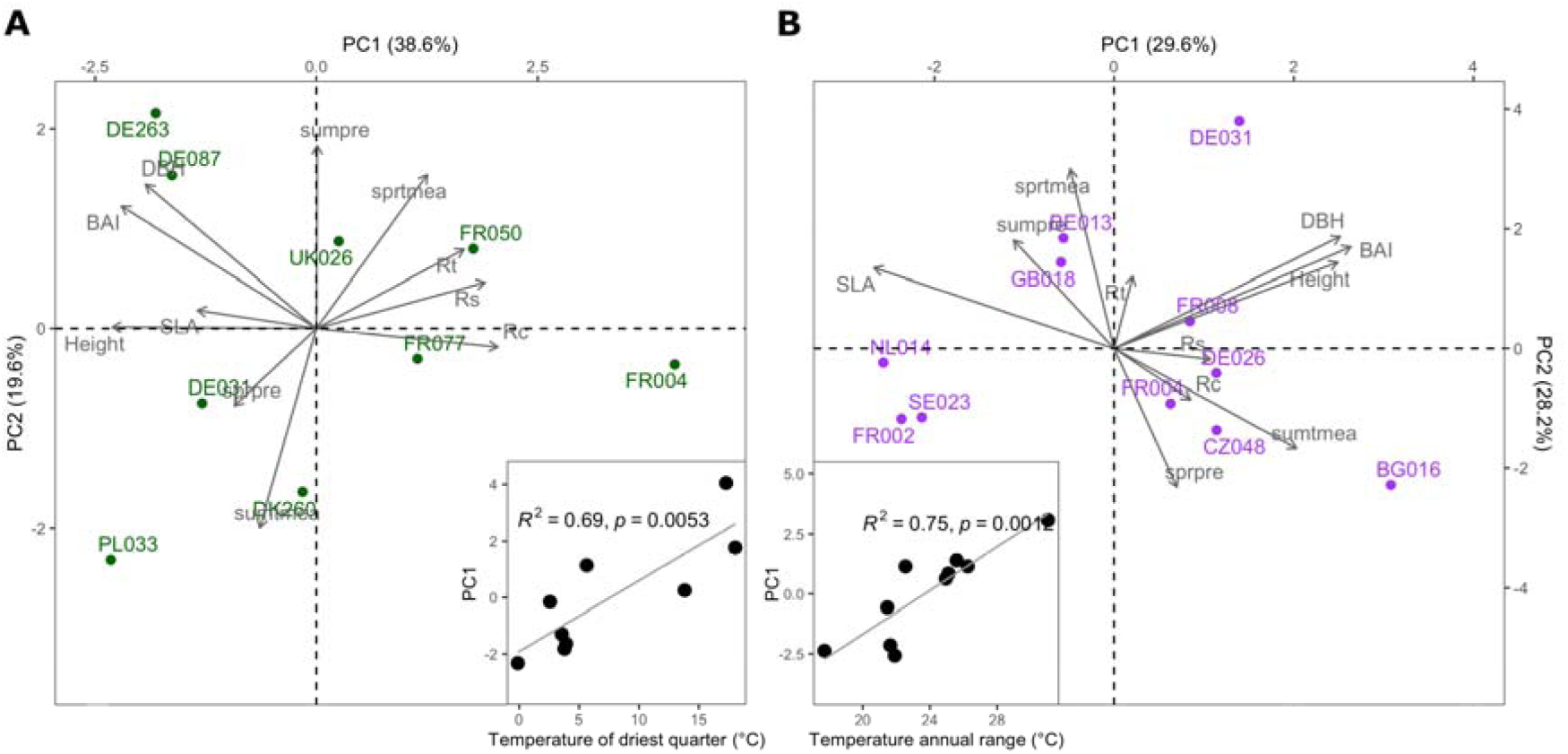
Principal component analyses of the provenance traits (only G effects) for oak **(A) and beech (B).** Inset plots show the regression of PC1 scores on climatic conditions of the provenances.

For beech, PC1 (29.6% of variance explained) scores were positively related to DBH, BAI, height, Rs, sumtmea and sprpre, but negatively related to SLA, sumpre and sprtmea (Figure 5B). The PC2 scores were positively correlated with height, BAI, DBH, Rt, sprtmea, sumpre and SLA, and negatively with sumtmea, sprpre, and Rc. The variance explained was more balanced between the two axes (PC2: 28.2%) than in oak. However, PC1 scores showed a significant relationship (linear regression) with climate of the provenance, specifically a positive relationship with temperature annual range (Figure 5B, inset plot), but also with mean diurnal range (R^2^= 0.63, *p*-value<0.01) and temperature seasonality (R^2^=0.54, *p*-value <0.05). As in the case of oak, no significant pattern was observed for PC2.

### Plasticity of multi-traits

When pooling trait provenance means from all common gardens (G × E effects), we observed a clear clustering by common gardens (Figure 6). The first two axes explained 64.6% (42.3% and 22.3%) and 53.9% (29.7% and 24.2%) for oak and beech, respectively. In oak, PC1 was positively correlated with DBH, BAI, height, sumtmea, sprtmea, Rs, Rt, and SLA, and negatively with sprpre and sumpre. PC2 was positively correlated with sprpre, sumpre, Rc, BAI, DBH, height, and sumtmea, and negatively with SLA, sprtmea and Rs. In beech, there was no trait negatively correlated with PC1 while sumpre, DBH, BAI, height and SLA were positively correlated. PC2 was positively correlated with sprtmea, Rt, Rs, SLA, sprpre and sumpre and negatively correlated with DBH, BAI, height, Rc and sumtmea.

**Figure 6.**
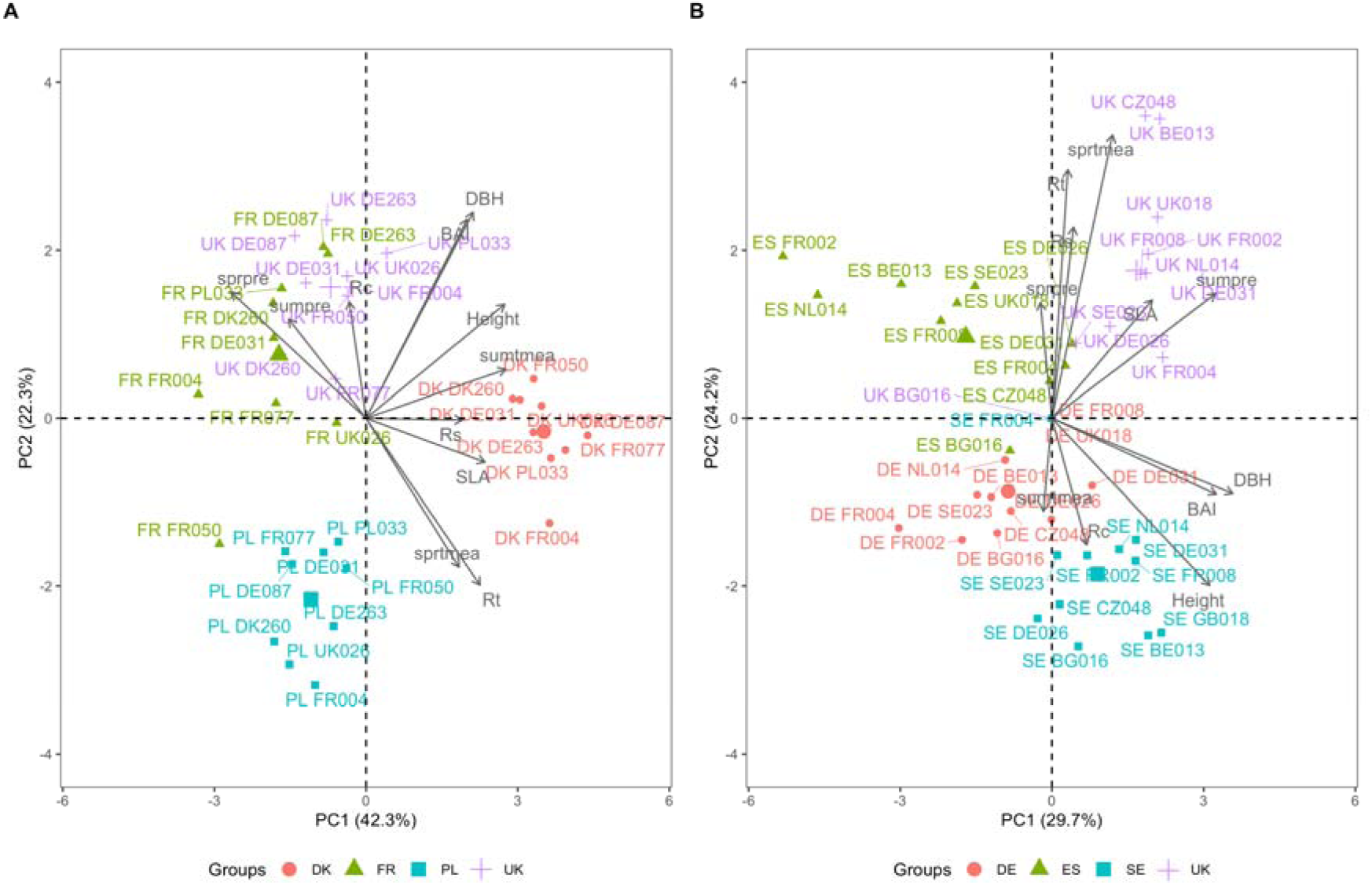
Principal component analyses of the phenotypic plasticity on multi-trait phenotypes across common gardens for oak (A) and beech (B). Larger symbols represent the common garden mean.

The maximum direction of changes (angle) was correlated with temperatures from the origin in both species, with provenances from warm areas showing greater ability to change the multi-trait phenotype (Figure S4). Specifically, the angle for oak was significantly positively related to temperature of the warm quarter and month. For beech, it was positively related to temperature of the coldest month and quarter, as well as annual temperature, but negatively related to temperature seasonality. Finally, the maximum magnitude of change (distance) was also positively correlated with temperature of the driest quarter for oak and negatively with temperature seasonality for beech.

## DISCUSSION

We analysed functional trait variation among provenances of sessile oak and European beech in common garden experiments across Europe to test whether growth- and morphological- related trait differences are due to genetic differences (provenance, G), environment (common garden, E), or their interaction (G × E). We interpret these factors as pure genetic effects (G), phenotypic plasticity (E), and genetically based plasticity (G × E, De Jong 1990; Vitasse *et al.,* 2013). Overall, our results indicate that variation in the studied traits of oak and beech is markedly driven by phenotypic plasticity although natural selection and local adaptation processes may still play an important secondary role. At the individual trait level, sessile oak showed evidence for natural selection as the driving force behind both genetic and plastic causes of phenotypic variation. In contrast, European beech seemed more prone to respond to environmental changes through plasticity with lighter indication for genetic effects and less signs of genetically based phenotypic plasticity compared to oak. The results of the multi-trait phenotypes, however, suggested genetically driven differences along a resource-use gradient governed by temperature-related conditions in both species. The plasticity of multi-trait phenotypes also reflects the capacity of all provenances to adjust to new conditions by optimizing the multi-trait phenotype along a resource-use gradient.

### Phenotypic plasticity dominates the response to different environments

The majority of phenotypic variation observed in our study was explained by the environmental conditions of the common gardens. Previous studies have also reported a high phenotypic plasticity in traits of juvenile oak and beech (Vitasse *et al*., 2010; Sáenz-Romero *et al*., 2017, 2019; GáratelEscamilla *et al*., 2019; Muffler *et al*., 2021; Schmeddes *et al*., 2024). Although our study design has limitations (four common gardens per species, covering only a part of the species’ distribution range but a large portion of their climatic niche), many of the oak traits followed temperature gradients of the common gardens whereas many of the beech traits were better correlated with precipitation of the common gardens. These patterns are in line with recent literature reporting European beech as a drought-sensitive species (Hacket-Pain *et al*., 2016; Dorado-Liñán *et al*., 2019), explaining its recent dieback in many parts of Central Europe due to extreme summer droughts since 2018 (Schuldt *et al*., 2020). In contrast, sessile oak has been described as a drought-tolerant (e.g., Martínez-Sancho *et al.,* 2017) and usually more temperature-limited species (but see Bose *et al.,* 2021).

Some authors relate the plasticity described above to a passive mechanism (and therefore inevitable plasticity) due to biochemical or physiological constraints (Pigliucci *et al*., 2006). Other studies, however, claim that such responses result from an "active" process, thus, considering phenotypic plasticity as an important mechanism shaping the response to natural selection imposed by changing/different environments (De Jong, 2005). For oaks, most of the variation in phenotypic plasticity between provenances was explained by temperature of origin, implying a genetic basis of the plastic responses to different climate conditions. As formulated by De Jong, (1995, 2005), if phenotypic plasticity has a strong genetic basis, it could be considered a trait on which natural selection can act. This implies a key role of phenotypic plasticity in shaping phenotypic change by interacting with intrinsic evolutionary processes such as genetic adaptation, genetic drift or mutation. Here, we found that oaks from warmer provenances exhibited the greatest range of responses in height and recovery from extreme events, whereas oak provenances from colder environments appeared to have a reduced range of responses of secondary growth. This finding aligns with observations in shoot growth among provenances of cork oak (Ramirez-Valiente *et al*., 2010) and sessile and

English oaks (Bert *et al*., 2020). Recent studies reanalysing juvenile growth in oak common gardens (including the network studied here) also found a high resilience to drought through phenotypic plasticity in provenances from warmer origins but not the potential of provenances from warmer sites to grow taller in cooler environments (Mátyás, 2021). This difference between our results and those reported by literature could be explained by contrasting age-related strategies. Although the G × E interactions were also statistically significant in all traits of beech, we found only a few associations between climatic variables from the provenances and genetically based plasticity of traits suggesting a low selective pressure acting on phenotypic plasticity for this species. This weak selection on phenotypic plasticity is in line with other studies assessing growth and physiological traits of beech seedlings across precipitation gradients (Knutzen *et al*., 2015). Our study also highlights the prominent role of plasticity when assessed using multi-trait phenotypes. Provenances from both species acclimatized to the common garden conditions by adjusting a combination of traits. In addition, the plasticity of these multi-trait phenotypes was partly related to climate of origin. For oaks, we found that provenances from warmer regions exhibited the largest magnitude and direction of changes of phenotypes whereas it was not only temperature but also its seasonality that determined these changes for beech. These results are consistent with our observations of plasticity in individual traits. A global analysis of phenotypic plasticity also observed an increase in phenotypic plasticity in plant size towards warm areas (Stotz *et al.,* 2021, but see Arnold *et al*., 2019). Further studies, incorporating a wider variety of traits and spectra and deciphering the genetic basis of trait integration, are needed to comprehensively assess the overall strategies of the species and their responses to evolutionary forces.

### Climate as a major driver of provenance differentiation

In addition to phenotypic plasticity, genetically driven differences were also found in the studied traits and multi-trait phenotypes, with greater evidence for climate-induced selective pressures for oak than for beech. For oak, among-provenance differences in secondary growth, sensitivity to climate, and resilience to extreme drought were related to clines in temperature and precipitation. Several other studies with oaks revealed genetic differentiation in growth, phenology (Vitasse *et al*., 2009; Kremer *et al*., 2010; Sáenz-Romero *et al*., 2017, 2019) and tree-ring-related traits (Bert *et al*., 2020). Studies assessing resilience to extreme drought in conifers also observed genetic adaptation (Depardieu *et al*., 2020; Zas *et al*., 2020; Martínez-Sancho *et al*., 2021), as illustrated in this study. Kremer (2016) and Kremer & Hipp (2020) highlighted distinct evolutionary and genetic characteristics inherent to the entire *Quercus* genus including high genetic diversity, elevated rates of ecological divergence, and a propensity for hybridization and introgression that may favour adaptation as a main evolutionary driver despite extensive gene flow. This was supported by e.g. genome-environment associations (GEAs) that revealed clear molecular genetic signatures of adaptation to climate and soil conditions in white oaks on a regional scale (Rellstab *et al.,* 2016b) and common garden experiments that highlighted the important role of introgression in climate adaptation (Leroy *et al*., 2020).

The results of our climate-transfer correlations, however, mostly related local adaptation processes to annual precipitation. Such associations between local adaptation patterns with precipitation resembled the previous reported associations between survival rate and height of seedlings with dryness (Sáenz-Romero *et al*., 2017). Interestingly, our results also highlight the signs of local adaptation in growth sensitivity to summer temperatures. This result is important since the genetic component of growth climate sensitivity is usually overlooked when analysing, inferring and projecting climatic sensitivity applying a space-for-time approach using large networks of tree-ring series (e.g., Martinez Del Castillo *et al*., 2022). Such genetically based growth sensitivity might also be relevant and necessary to integrate into, for instance, assisted gene flow frameworks.

We observed a significant effect of beech provenance in nine out of the eleven studied traits, although the clinal patterns relative to climate of origin were relatively weak. Such reduced signs of climate adaptation in beech traits has also been previously reported in early life-history traits (Muffler *et al*., 2021; Schmeddes *et al*., 2024), physiological traits (Knutzen *et al*., 2015), and phenology (Vitasse *et al*., 2010; but see Robson *et al*., 2013). Although no significant correlations between height (G effects) and the climate of origin were observed, we found evidence of local adaptation to precipitation in the coldest quarter when analysing climate-transfer distance. Previous studies reported a significant association between tree height and the potential evapotranspiration of the provenances (Gárate-Escamilla *et al*., 2019), further supporting our findings related to hydric requirements. In our study, SLA showed a correlation with temperature-related climate variables at provenance origin. These results aligned well with those from the climate-transfer distance since SLA showed signs of local adaptation to temperature of the warmest quarter. Although SLA has been primarily associated with precipitation clines (e.g., Rose *et al*., 2009; Robson *et al*., 2012), other studies also found significant imprint of temperature of origin when testing provenances from an elevation gradient (Bresson *et al*., 2011).

Interestingly, we observed an exponential relationship with mean temperature of the wettest quarter, rather than quadratic, in the climate-transfer distance of sensitivity of growth to spring temperature (sprtmea) of beech explaining a large part of the variance (52%). This suggests that when plants are transferred towards warmer areas, growth sensitivity to spring temperature is low. In contrast, plants move to colder environments exhibit increased signs of limiting temperature requirements (positive correlations). These results indicate that provenances are adapted to minimum temperature thresholds required for cambium reactivation in spring, which are not met when they are transferred to colder environments (Rathgeber *et al*., 2016), and therefore, this adaptation is more strongly expressed. When assessing drivers of among-provenance differences, recovery after extreme drought (Rc) emerged as being strongly linked to precipitation clines, suggesting that provenances from wetter origins recovered less from extreme droughts. This association was also found in common garden studies of white spruce (Depardieu *et al*., 2020). However, we could not attribute this pattern to local adaptation processes analysed with climate-transfer distance. Instead, we found a significant pattern of local adaptation in resistance (Rt, with an opposite pattern in Rc) related to isothermality. Beech trees may have adapted to their thermal range conditions, either optimizing their growth cycles, cambial activity, and stomatal regulation in relatively stable (high-isothermality) environments or developing traits like increased stress tolerance in more variable (low-isothermality) conditions to cope with greater temperature extremes. Variations in these ranges might lead to maladaptation. Not surprisingly, Rc displayed the opposite pattern, as there is a close relationship between the two metrics. However, the maximum (minimum) rates of Rt and Rc were found at negative climate- transfer distance values, which suggests adaptation lags as a result of an environmental change or gene flow (RamírezlValiente *et al*., 2022). It is also important mentioning that biotic conditions (i.e., competition) explained a significant part of the variance of the models for both species, specifically those models related to growth traits. This result was expected as numerous studies have reported a negative relation between growth and competition (Coomes & Allen, 2007; Pretzsch *et al*., 2013). This significant effect of competition on resilience components in beech is in line with previous research (Castagneri *et al*., 2022). Thus, by accounting for competition, we were able to better characterize growth conditions and obtain more refined estimates of provenance and site effects.

Interestingly, we observed temperature-related clines for the multi-trait phenotypes in both species, showing that selection acts simultaneously on multiple traits that are potentially correlated with each other, ultimately affecting the whole phenotype (Pigliucci *et al*., 2006; Díaz *et al*., 2016; MartínezlVilalta *et al*., 2023). The multi-trait phenotypes of both species follow a gradient of resource-use strategy according to Reich (2014), showing a clear growth- drought resistance trade-off. Individuals originating from cold environments displayed higher rates of both primary and secondary growth. However, the two species diverged in the strategy exhibited at the opposite extremes of the gradient. Individuals of sessile oak from warmer provenances coped better with extreme events, whereas those of beech showed higher regulation of SLA. Previous studies have reported a high regulation of beech leaf traits in response to drought conditions (Pflug *et al*., 2018), suggesting that this is the primary control for the drought strategy of a species. Studies assessing the multi-trait phenotype of a different *Fagaceae* species also found this climate-controlled resource-use gradient on trait coordination (Rosas *et al*., 2019; SolélMedina *et al*., 2022). Our study also highlights the potential of dendrochronologically derived traits for addressing important evolutionary research questions (Housset *et al*., 2018). Further studies assessing the capacity of a species to resist climate change through trait variation and its plasticity may consider incorporating gene expression analyses with tree-ring-related, physiological, hydraulic, or belowground- related traits to obtain a comprehensive understanding of species adjustments.

### Implications for the evolutionary strategies of species

The outcome of our study suggests that environmentally and genetically based phenotypic plasticity is a key process driving trait variability of sessile oak and European beech across their natural distribution range. Previous studies have classified both species as adaptive generalists (species prevalently acclimating to new conditions through phenotypic plasticity) rather than adaptive specialists (species prevalently adapting to new conditions through genetic adaptation) (see Leites & Benito Garzón 2023). In our study, this generalist strategy is less obvious in the case of oak, since genetic processes play a role in shaping single trait patterns besides phenotypic plasticity. In contrast, this generalist strategy is more pronounced in beech since we found little evidence of climate variation shaping trait variability. However, we observed a clinal gradient in the multi-trait phenotype of the species suggesting that climate adaptation processes might not be as evident in single traits but is revealed when integrating multiple traits, highlighting its necessity in quantitative genetics. The role of phenotypic plasticity in evolution is still controversial, with debates centred on whether plasticity reduces the efficiency of natural selection or creates new opportunities for selection to act on (Ghalambor *et al*., 2007; Anderson *et al*., 2021). The issue is especially relevant in canopy-forming, long-lived forest trees with long generation time and heavy seeds, where parental trees compete with their offspring for space and light (Petit & Hampe 2006). While the long time needed to reach fecundity requires a high degree of phenotypic plasticity in the parental trees, it might lead to a substantial maladaptation in the offspring generation when environmental changes are fast. For European beech, the reduced genetically based responses and its reliance on plastic responses to cope with novel conditions might be a safe strategy at first, but it also threatens its capacity to survive once the limits of plasticity are surpassed, leading to potentially large-scale dieback, as recently observed after the extreme hot drought in 2018 in Europe (Schuldt et al 2020). In contrast, the fact that there are signs of natural selection in oak responses might prevent this species from abrupt decline as climate continues to warm and extreme drought to occur. Our results are crucial for assisted migration and climate-focused seed programs since they usually seek provenances adapted to warmer/drier conditions as future forest resources. Our results show that this strategy is not false, but increased attention could be given to provenances showing high degrees of plasticity with respect to climate. Because it is impossible to disentangle plasticity due to local microclimatic differences and adaptive genetic diversity in natural provenances, common garden experiments remain the gold standard to quantify the relative contribution of plasticity and adaptation to trait variation and fitness. Further research should consider incorporating an integrative set of traits to understand the overall ability of tree species and provenances to cope with forthcoming climate change effects.

## ACKNOWLEDGEMENTS

This project was fully funded by the SwissForestLab (“Common-Ring project”). EM-S was partly funded by the project RYC2021-035078-I from the Spanish Ministry of Science and Innovation. JCM was supported by the María Zambrano fellowship (reference: RMZ-21- DJVHZ3-7-90M8PD) from the Spanish Ministry of Universities with European Union-Next Generation EU funds. We are also thankful to Sixtine Guinard, Joanna Reim and Isa van Lidth de Jeude for their support during field campaigns and to Gustavo López for his help in the common garden selection.

## AUTHOR CONTRIBUTION

EM-S, together with CR and YV, let the design of the study with contributions from PF, MBG and CB. JCM, YV and PF in collaboration with the managers of the common gardens (DJC, JC, EDK, AD, JKH, MK, ML, LR, and RW) performed the field sampling. AV and MN performed the tree-ring measurements. EM-S revised the measurements, homogenized the dataset, and performed the statistical analyses. EM-S, with help from CR and YV, led the writing of the manuscript. All the authors have revised and contributed to the final version of the manuscript.

