## supplementary for "Genetic and plastic effects on trait variability in two major tree species: insights from common garden experiments across Europe"

**Supplementary Information**

**Table S1.** Location of the studied common gardens.

**Table S2.** Geographic origin of the oak and beech provenances.

**Table S3.** Output of the linear mixed models.

**Table S4.** Statistics of the fitted quadratic functions between trait variation and climatic transfer distances.

**Figure S1.** Climate envelope based on mean annual temperature and annual precipitation for oak and beech.

**Figure S2.** Temporal evolution of different water balances for all the studied sites.

**Figure S3.** Pearson correlations between least-square means and common garden climate conditions for the period 1987-2021.

**Figure S4.** Pearson correlation between trait direction (angle) and magnitude (distance) and climatic conditions of seed origin.

**Table S1. Location of the studied common gardens.**

| Species | Country | Site | Country code | Latitude (DD) | Longitude (DD) |
| --- | --- | --- | --- | --- | --- |
| Oak | France | Vierzon | FR | 47.26 | 2.13 |
| Oak | United Kingdom | Arden | UK | 52.33 | -1.48 |
| Oak | Poland | Kornik | PL | 52.24 | 17.08 |
| Oak | Denmark | Valby hegn | DK | 56.05 | 12.22 |
| Beech | Spain | Pazuengos | ES | 42.30 | -2.92 |
| Beech | Germany | Liliental | DE | 48.07 | 7.67 |
| Beech | United Kingdom | Oxford | UK | 51.63 | -1.18 |
| Beech | Sweden | Trolleholm | SE | 55.93 | 13.32 |

**Table S2. Geographic origin of the oak and beech provenances**. N sites refers to the number of experimental sites where the provenance was planted.

| Species | ID | Country | Latitude (DD) | Longitude (DD) | N sites |
| --- | --- | --- | --- | --- | --- |
| Oak | DK260 | Denmark | 56.07 | 9.6 | 4 |
| Oak | FR004 | France | 47.81 | 0.39 | 4 |
| Oak | FR050 | France | 46.6 | 0.18 | 4 |
| Oak | FR077 | France | 47.76 | 5.49 | 4 |
| Oak | DE031 | Germany | 52.83 | 10.32 | 4 |
| Oak | DE087 | Germany | 49.36 | 7.87 | 4 |
| Oak | DE263 | Germany | 53.71 | 9.76 | 4 |
| Oak | PL033 | Poland | 51.18 | 17.93 | 4 |
| Oak | UK026 | United Kingdom | 51.95 | -2.45 | 4 |
| Beech | BE013 | Belgium | 50.83 | 4.42 | 4 |
| Beech | BG016 | Bulgaria | 41.57 | 23.73 | 3 |
| Beech | CZ048 | Czech Republic | 50.8 | 15.23 | 3 |
| Beech | FR002 | France | 49.53 | 0.77 | 4 |
| Beech | FR004 | France | 44.15 | 2.58 | 3 |
| Beech | FR008 | France | 42.92 | 2.32 | 3 |
| Beech | DE026 | Germany | 53.65 | 10.67 | 4 |
| Beech | DE031 | Germany | 48.47 | 9.45 | 4 |
| Beech | NL014 | Netherlands | 51.93 | 6.73 | 4 |
| Beech | SE023 | Sweden | 55.57 | 13.2 | 4 |
| Beech | UK018 | United Kingdom | 51.72 | -2 | 3 |

**Table S3. Statistical outputs of the linear mixed-effects models.** Significance of the fixed-effects; site (common garden), provenance, site×provenance and competition with *F* for traits fitted with linear-mixed models. Significant effects are shown in bold *(p*<0.05).

| **Oak** | SITE (E) |  | PROVENANCE (G) | | SITE x PROVENANCE (GxE) | | COMPETITION | |
| --- | --- | --- | --- | --- | --- | --- | --- | --- |
|  | F | *P* | F | *P* | F | *P* | F | *P* |
| DBH | 17.414 | **<0.001** | 3.165 | **0.002** | 4.155 | **<0.001** | 300.293 | **<0.001** |
| Height | 8.694 | **0.002** | 8.641 | **<0.001** | 5.865 | **<0.001** | 123.278 | **<0.001** |
| SLA | 41.852 | **<0.001** | 1.582 | 0.128 | 3.010 | **<0.001** | 8.998 | **0.003** |
| BAI | 12.011 | **<0.001** | 46.713 | **<0.001** | 25.856 | **<0.001** | 1847.565 | **<0.001** |
| sprpre | 48.212 | **<0.001** | 0.729 | 0.666 | 2.106 | **0.002** | 2.634 | 0.105 |
| sumpre | 12.286 | **0.004** | 1.027 | 0.415 | 2.155 | **0.001** | 3.829 | 0.051 |
| sprtmea | 16.116 | **0.002** | 2.189 | **0.027** | 2.469 | **0.000** | 11.922 | **0.001** |
| sumtmea | 92.109 | **0.011** | 2.212 | **<0.001** | 2.388 | **0.000** | 0.164 | 0.686 |
| Rs | 9.906 | **0.001** | 1.761 | 0.082 | 3.288 | **<0.001** | 0.408 | 0.523 |
| Rt | 5.524 | **0.021** | 1.295 | 0.244 | 1.907 | **0.006** | 1.748 | 0.187 |
| Rc | 67.960 | **<0.001** | 1.592 | 0.125 | 4.224 | **<0.001** | 6.841 | **0.009** |
| **Beech** |  |  |  |  |  |  |  |  |
| SLA | 12.373 | **0.015** | 5.982 | **<0.001** | 4.835 | **<0.001** | 5.626 | **0.018** |
| DBH | 36.212 | **<0.001** | 9.243 | **<0.001** | 5.177 | **<0.001** | 257.231 | **<0.001** |
| Height | 11.892 | **0.015** | 11.354 | **<0.001** | 8.200 | **<0.001** | 63.020 | **<0.001** |
| BAI | 2.766 | 0.157 | 52.355 | **<0.001** | 36.917 | **<0.001** | 1909.774 | **<0.001** |
| sprpre | 0.620 | 0.633 | 1.067 | 0.386 | 1.955 | **0.004** | 14.508 | **<0.001** |
| sumpre | 4.537 | 0.092 | 0.623 | 0.795 | 3.309 | **<0.001** | 11.078 | **<0.001** |
| sprtmea | 7.571 | **0.035** | 2.138 | **0.020** | 1.771 | **0.013** | 8.781 | **0.003** |
| sumtmea | 1.356 | 0.382 | 2.697 | **0.003** | 2.638 | **<0.001** | 1.580 | 0.209 |
| Rs | 4.258 | 0.089 | 2.995 | **0.001** | 3.185 | **<0.001** | 23.262 | **<0.001** |
| Rt | 61.510 | **0.010** | 2.934 | **0.001** | 2.412 | **<0.001** | 15.951 | **<0.001** |
| Rc | 83.625 | **<0.001** | 2.209 | **0.016** | 3.321 | **<0.001** | 0.002 | 0.967 |

**Table S4. Statistics of the fitted quadratic functions between trait variation and climatic transfer distances.** *F* values from an ANOVA test and ∆BIC refers to the comparison of the quadratic model with a linear model.

| SPECIES | TRAIT | BIOCLIM | R2 | *p*-value | *F* | ∆BIC |
| --- | --- | --- | --- | --- | --- | --- |
| Oak | DBH | bio12 | 0.41 | 0.000 | 0.001 | 9.26 |
| Oak | DBH | bio3 | 0.21 | 0.019 | 0.008 | 4.32 |
| Oak | Height | bio12 | 0.30 | 0.003 | 0.001 | 9.30 |
| Oak | SLA | bio12 | 0.51 | 0.000 | 0.001 | 8.08 |
| Oak | BAI | bio12 | 0.33 | 0.001 | 0.009 | 4.00 |
| Oak | BAI | bio3 | 0.17 | 0.045 | 0.016 | 2.80 |
| Oak | sumpre | bio2 | 0.22 | 0.018 | 0.023 | 2.13 |
| Oak | sumtmea | bio12 | 0.44 | 0.000 | 0.000 | 16.63 |
| Oak | sumtmea | bio3 | 0.35 | 0.001 | 0.012 | 3.37 |
| Oak | Rc | bio15 | 0.38 | 0.000 | 0.021 | 2.28 |
| Beech | Height | bio19 | 0.21 | 0.020 | 0.017 | 2.78 |
| Beech | SLA | bio10 | 0.31 | 0.002 | 0.002 | 7.35 |
| Beech | SLA | bio5 | 0.28 | 0.005 | 0.003 | 6.03 |
| Beech | sprtmea | bio8 | 0.52 | 0.000 | 0.020 | 2.42 |
| Beech | Rt | bio3 | 0.32 | 0.002 | 0.005 | 5.28 |
| Beech | Rc | bio3 | 0.35 | 0.001 | 0.002 | 7.52 |

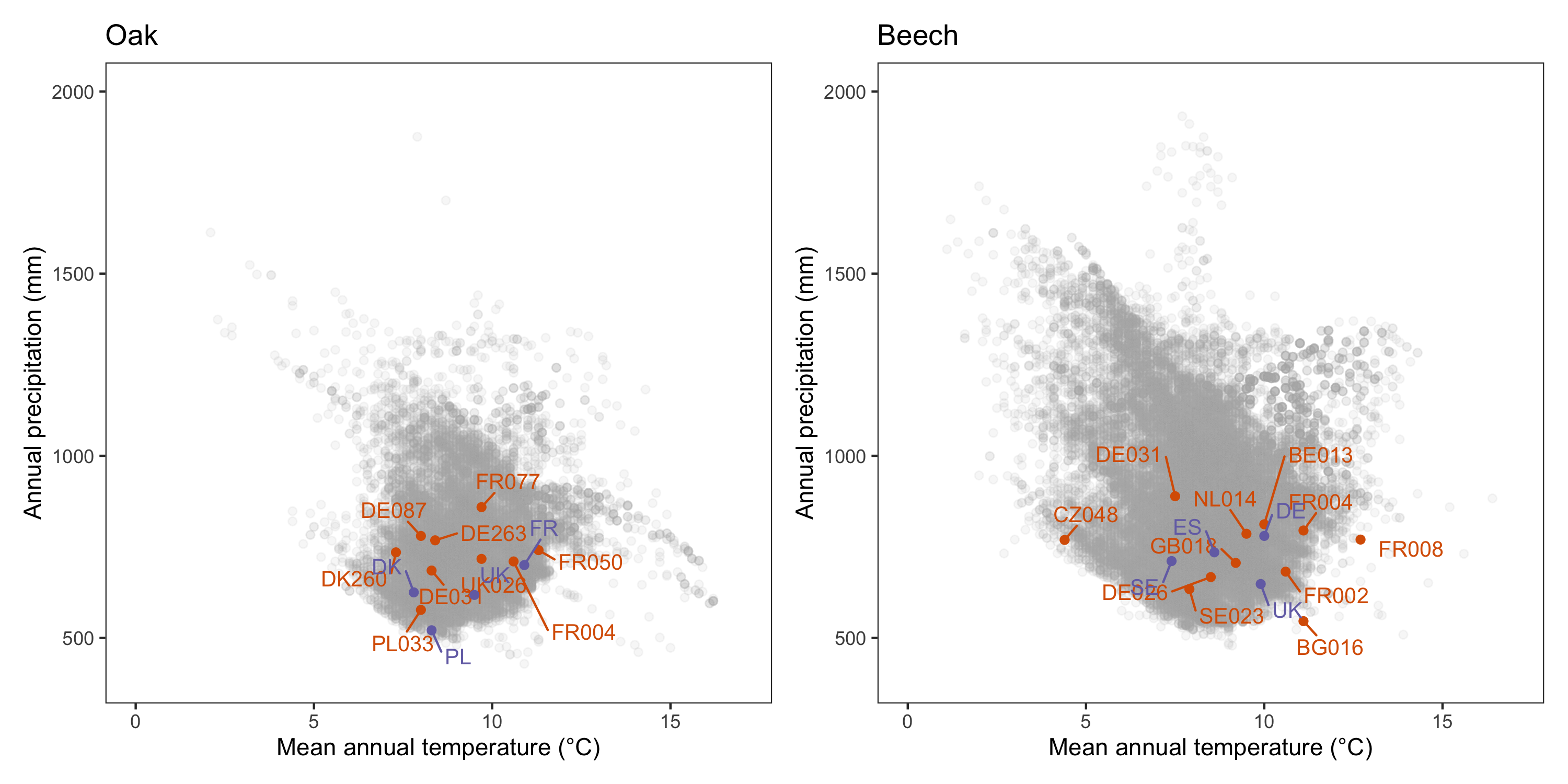

**Figure S1. Climate envelope based on mean annual temperature and precipitation for oak and beech.** Grey points represent species occurrences from the European distribution obtained from Mauri *et al.* (2017). Orange and purple dots correspond to the climatic position of the selected provenances and common gardens, respectively. All data were extracted from the Worldclim database for the period 1970-2000.

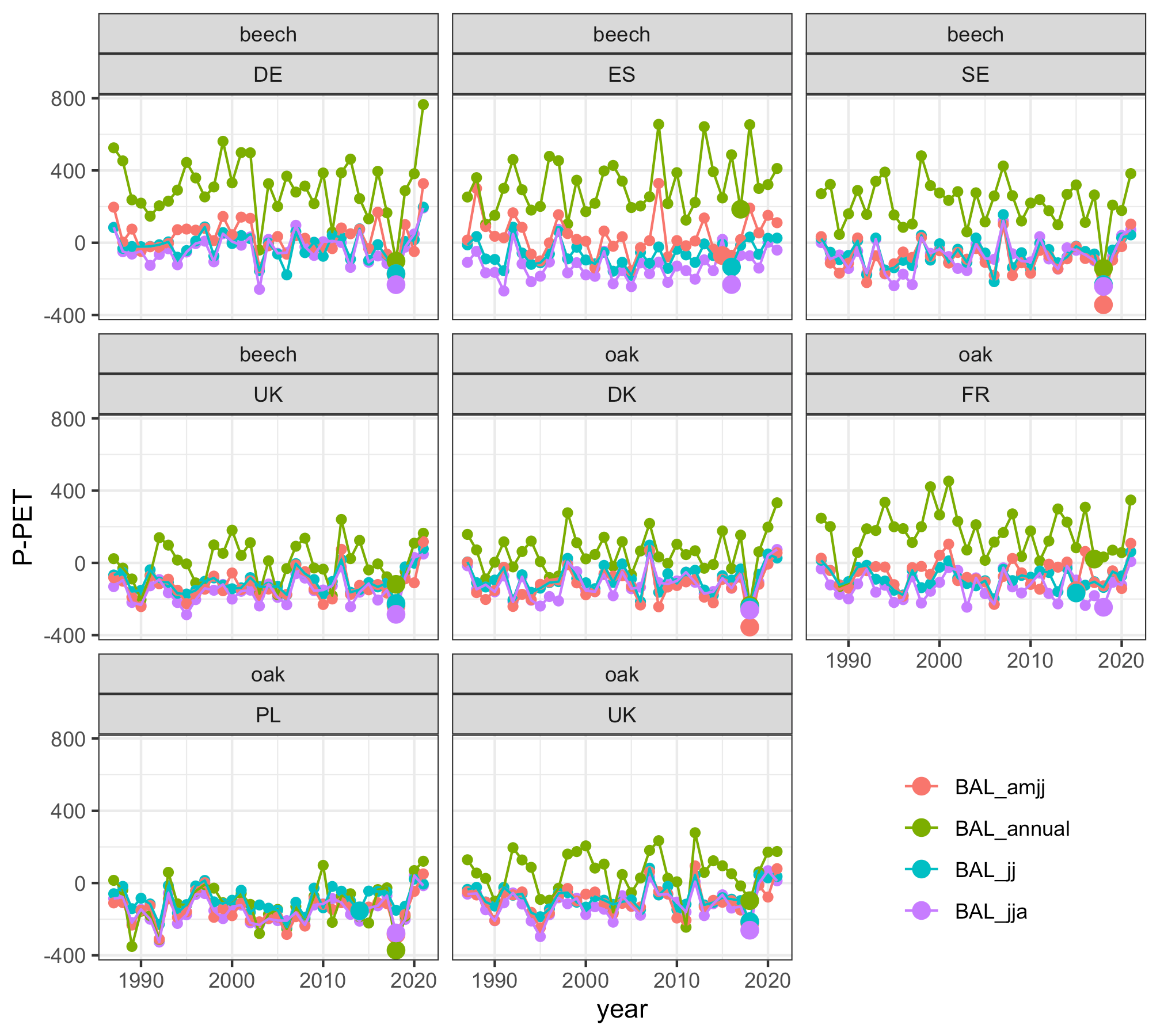

**Figure S2. Temporal development of climatic water balance for all the studied sites computed over different time windows.** BAL_amjj_ = water balance calculated from April to July; BAL_annual_ = annual water balance; BAL_jj_ = water balance calculated from June to July; BAL_jja_ = water balance calculated from June to August. Larger symbols represent the year with the lowest water balance.

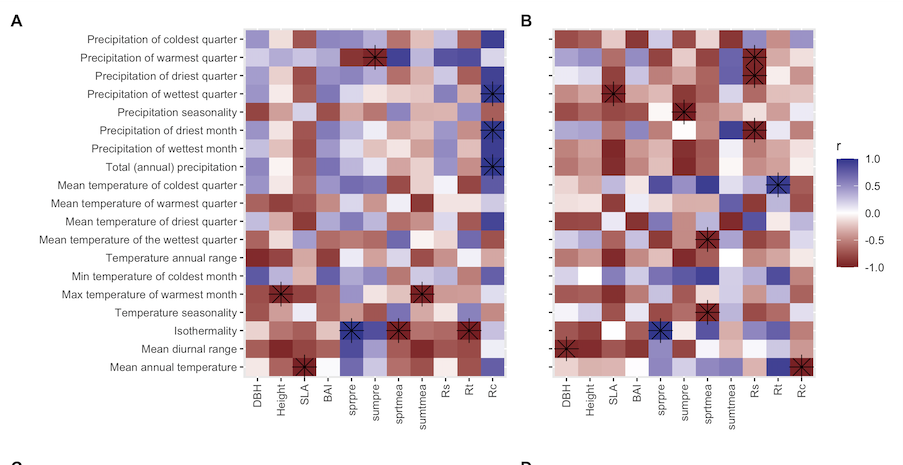

**Figure S3. Pearson correlations between least-square means and common garden climate conditions for the period 1987-2021 for oak (A) and beech (B).** Asterisks indicate significant correlations at *p*< 0.05 level.

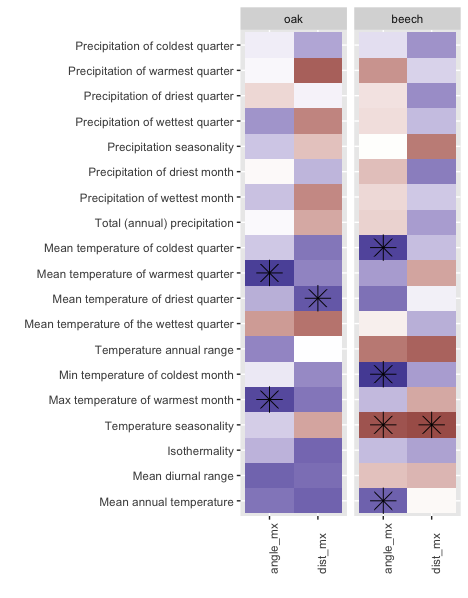

**Figure S4. Pearson correlation between trait direction (angle_mx) or magnitude (dst_mx) and climatic conditions at seed origin.** Asterisks indicate significant correlations at *p*< 0.05 level.
